# Punctuated mutagenesis promotes multi-step evolutionary adaptation in human cancers

**DOI:** 10.1101/2025.06.30.662384

**Authors:** Christopher Graser, Wenbo Wu, Cole Christini, Mia Petljak, Franziska Michor

**Affiliations:** Department of Data Science, Dana-Farber Cancer Institute, Boston, MA, USA; Department of Biostatistics, Harvard T.H. Chan School of Public Health, Boston, MA, USA; Department of Stem Cell and Regenerative Biology, Harvard University, Cambridge, MA, USA; School of Computer Science, Carnegie Mellon University; Department of Pathology, New York University School of Medicine, New York, NY, USA; Laura and Isaac Perlmutter Cancer Center, New York University, New York, NY, USA; Center for Cancer Evolution, Dana-Farber Cancer Institute, Boston, MA, USA; The Eli and Edythe L. Broad Institute, Cambridge, MA, USA; Ludwig Center at Harvard, Harvard Medical School, Boston, MA, USA

## Abstract

The rate of acquisition of genomic changes in cancer has been the topic of much discussion, with several recent investigations finding evidence of punctuated evolution instead of gradual accumulation of such changes. Despite forays into the description and quantification of these punctuated events, the effects of such changes on subsequent cancer evolution remain incompletely understood. Here we investigate how non-gradual mutagenesis affects the ability of tumor cells to acquire and retain fitness-enhancing adaptations. We find that punctuated mutagenesis significantly facilitates adaptation in scenarios where adaptation requires crossing a fitness valley, i.e. when multiple mutations are required which individually are maladaptive but jointly confer a fitness advantage. By increasing the probability that multiple mutations occur in close succession, punctuation increases the chance that mutants in a fitness valley mutate further to reach a fitness peak before going extinct. Analyzing data from The Cancer Genome Atlas, we find that tumors with signatures of APOBEC mutagenesis, which has been shown to proceed in episodic bursts, exhibit patterns consistent with higher rates of crossing fitness valleys. Lastly, we characterize how the interplay between this enhanced ability to cross fitness valleys and adaptation-limiting effects of clonal interference affects overall adaptability in complex fitness landscapes.

## Introduction

Tumors evolve via the acquisition of randomly arising genetic and/or epigenetic alterations and their selection^1^. The rate at which tumor cells accumulate such genomic changes to produce potentially adaptive innovation is thus an important determinant of the capacity of tumor cells to adapt to diverse selection pressures^2,3^, affecting their propensity to progress locally, to metastasize, or to evolve resistance to therapeutic intervention. Several methods to quantify mutation rates have been developed^4–6^ and proxies for mutation rates such as tumor mutation burden often feature in prediction models of therapeutic outcomes^7,8^.

Most existing methods deal with constant mutation rates^6,9^, assuming gradual evolutionary change over time. However, accumulating evidence suggests that mutagenesis in cancer is fluctuating. Indeed, a recent study^10^ demonstrated in vitro that mutations associated with DNA-editing activity of APOBEC cytidine deaminases^11^ occur in episodic bursts, with more than 100-fold differences in the rate of APOBEC-associated mutations across otherwise identical cell culture replicates (Fig 1A). This episodicity observed in vitro aligned with patterns in previously published in vivo data investigating APOBEC mutagenesis in lung cancer^12,13^, and data from intestinal crypts^14^ later confirmed explicitly that this episodic pattern also occurs in patients. Other recent studies using genomic data to reconstruct evolutionary histories of tumors have found evidence of punctuated evolution across several cancer types^15–17^. These studies have demonstrated the existence of distinct phases of mutation bursts^17,18^ (Fig 1B), challenging the prevailing paradigm of gradual emergence of mutant lineages (Fig 1C). Various mutagenic processes such as chromothripsis^19,20^, chromoplexy^21^ and other drivers of chromosomal instability^22^ have been implicated in causing such punctuated patterns^23^.

**Figure 1:**
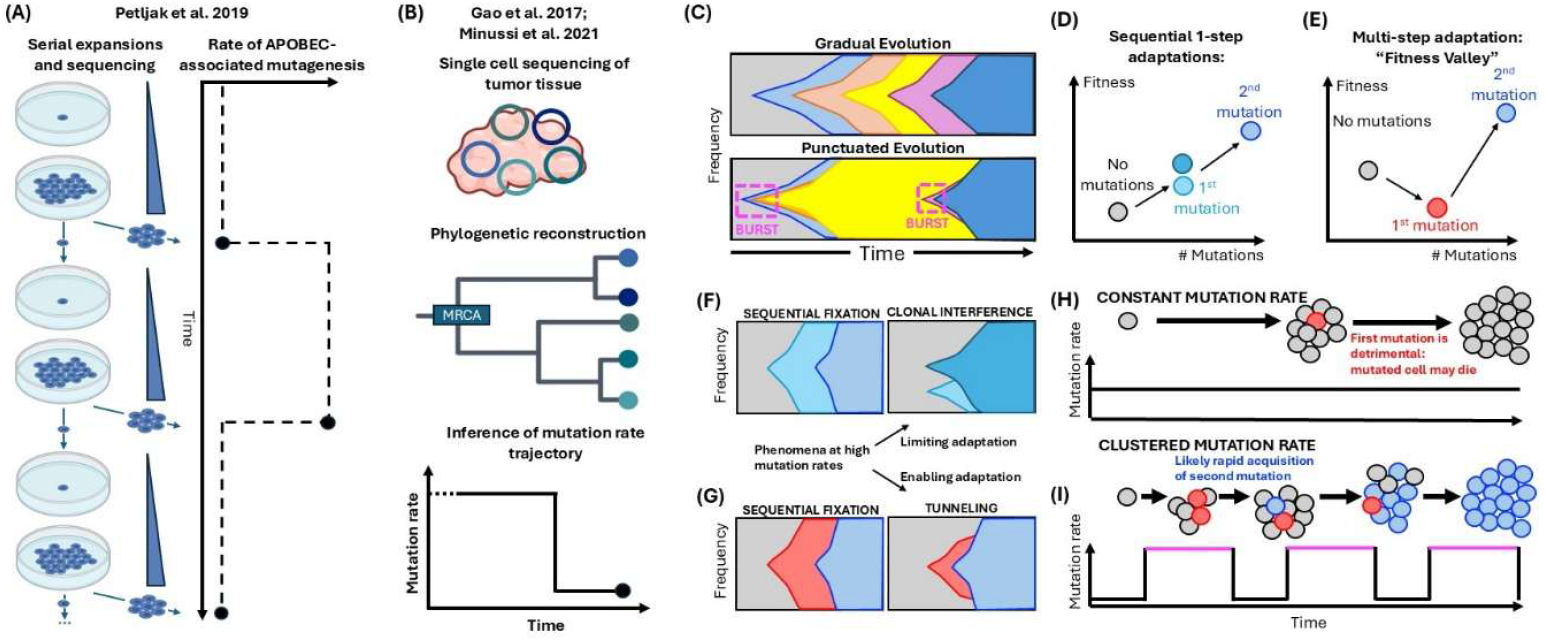
Population dynamics under gradual vs punctuated mutagenesis. **(A)** Evidence for punctuated APOBEC mutagenesis (Petljak et al. 2019) from successive cell culture expansions, seeded with single progenitors from the preceding expansion. Sequencing of the expanded populations revealed large fluctuations in APOBEC-associated mutagenic signatures SBS2 and SBS13. **(B)** Evidence for punctuated copy-number evolution (Gao et al. 2017, Minussi et al. 2021). Patterns in branch lengths of reconstructed phylogenetic trees reveal fluctuations in the rates of copy number alterations. Dynamics prior to most recent common ancestor (MRCA) are unidentifiable. **(C)** Gradual vs. punctuated evolution. In gradual evolution novelty emerges and spreads at a constant rate over time. In punctuated evolution novelty emerges and spreads during distinct burst phases. **(D)** Fitness schematic of two 1-step adaptations which each independently confer a fitness advantage. **(E)** Fitness schematic for a 2-step adaptation, in which carrying one mutation confers a fitness disadvantage but carrying two mutations confers a fitness advantage. **(F,G)** Schematic of possible evolutionary dynamics for two 1-step adaptations (F) and of modes of valley-crossing with or without prior fixation of the first mutant (G). Sequential fixation occurs at low mutation rates where emerging mutant lineages are likely to have fixated or gone extinct before the next mutation occurs. At higher mutation rates, the second mutation can occur in a multi-clonal population. **(H)** Sketch of failing two-step adaptation under a uniform (time-invariant) mutation rate. **(I)** Sketch of a successful two-step adaptation via stochastic tunneling under a punctuated mutation process with distinct clusters of high mutation rates.

How punctuated mutagenesis affects the evolutionary dynamics of tumor cell adaptation across complex fitness landscapes remains incompletely understood. Addressing this question with existing experimental data is inconclusive, as only short time horizons are observed (single expansions^10^; time to a most recent common ancestor^17,18^), and as fitness differences between observed cell lineages are incompletely characterized. Mathematical modelling of human cancer genomics data, however, can elucidate the dynamics of adaptation during punctuated tumor evolution. Here, we set out to systematically investigate the evolutionary consequences of punctuated mutation acquisition during tumorigenesis using mathematical modeling and analysis of genomic data from The Cancer Genome Atlas (TCGA).

## Results

### Temporal clustering facilitates multi-step adaptation

We set out to investigate how *temporal clustering* of mutagenic events into distinct episodes affects the ability of a cell population to achieve multi-step evolutionary adaptations. We considered two scenarios: tumor evolution via a single advantageous step, such as an oncogenic adaptation^24^ (“1-step adaptations”, Fig 1D), and accumulation of multiple mutations with synergistic fitness effects (“multi-step adaptations”, Fig 1E). Indeed, the two hit-hypothesis^25^ for tumor suppressor genes (TGSs) suggests that inactivation of one copy of a TSG can be inconsequential or maladaptive, while bi-allelic inactivation confers a selective advantage^26^. While some TSGs exhibit (context-dependent) haploinsufficiency^27–29^, synergistic fitness effects for successive (epigenetic^27,28,30^) alterations in the same gene remain the prominent feature of TSGs. Analogously, there are ample examples of successive alterations across different genes acting synergistically^31–33^. High mutation rates are known to limit the rate of retained 1-step adaptations per emerging 1-step adaptation due to clonal interference^34–36^ (Fig 1F), which has been highlighted as a potential consequence of punctuated cancer evolution^37^. The dynamics of multi-step adaptation have been studied extensively^26,38–44^, elucidating two modes of evolution (Fig 1G): ‘sequential fixation’ refers to the scenario in which the second mutation only emerges after cells harboring the first mutant have taken over the population, while ‘stochastic tunneling’ refers to situations in which the second mutation arises sooner. In large populations, mutants with a selective disadvantage become vanishingly unlikely to reach fixation, so that stochastic tunneling becomes the dominant mode of evolution for multi-step adaptations.

To investigate how the rate of successful stochastic tunneling events – enabling multistep adaptation – is affected by punctuated mutagenesis, we simulated selection dynamics of a population of cells which proliferate according to a Wright-Fisher process ^45,46^ and accumulate mutations according to a constant (Fig 1H) or temporally clustered (Fig 1I) rate. The Wright-Fisher process models evolution as successive, non-overlapping generations of constant size. Each new generation is populated by drawing with replacement from the cells in the previous generation, with probabilities proportional to their fitness (Fig 2A). Cells can accumulate mutations that change their fitness, i.e. increase their probability to survive to the next generation. We considered fitness landscapes in which two-step adaptations are the only way by which cells can increase their fitness, with fixed fitness ratios across subsequent two-step adaptations (Fig 2B,C). We also confirmed that the findings observed for the Wright-Fisher process, described below, are qualitatively consistent for expanding populations that proliferate according to a branching process model (Methods, Fig S1).

**Figure 2:**
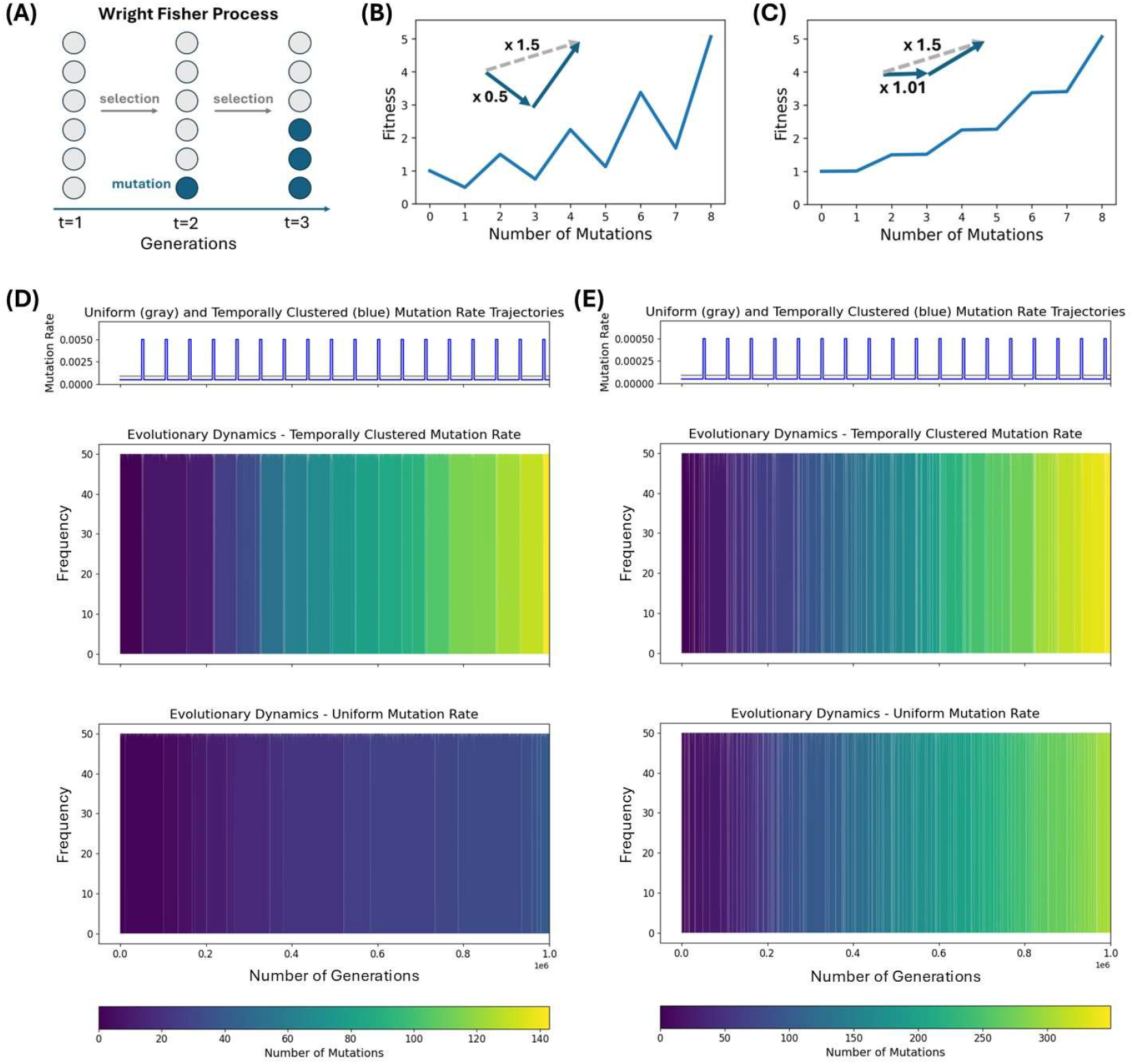
Simulation results: valley-crossing under uniform vs. temporally clustered mutation rates. **(A)** Schematic of a Wright Fisher Process. **(B)-(C)** Fitness landscapes used for the simulations in Panel (D) and (E) respectively. Mutations move a cell from left to right through the landscape. Each 2-step adaptation corresponds to a fitness increase by a factor of 1.5. Having an odd number of mutations comes at a multiplicative fitness disadvantage of 0.5 in (B) and an advantage of 1.01 in panel (D). **(D)-(E)** Heatmaps of simulation results for a Wright-Fisher process with 50 cells. The mutation rate trajectories in each panel are chosen such that the total expected number of mutations under the uniform trajectory is identical to that under the temporally clustered trajectory. Each column of pixels represents a population after a specific number of generations. Homogeneity in pixel-color per column thus indicates that cells with a given number of adaptations have fixated in the population, whereas persistent diversity of pixel colors (see Fig S1) indicates coexistence of lineages with different numbers of adaptations.

Interestingly, for two-step adaptations for which the intermediate mutants that carry only one of the two required mutations have a strong selective disadvantage (Fig 2B), we found that the population undergoes substantially (3.55 times) more two-step adaptations when accumulating mutations in a temporally clustered rather than in a uniform way (Fig 2D). This effect arises because intermediate mutants in “fitness valleys” have a high chance of going extinct before acquiring the next mutation. During mutation bursts, acquiring the next mutation in time before the disadvantageous mutant goes extinct becomes more likely. Similarly, intermediate mutants are more likely to emerge in a mutation burst. This temporal clustering of the likelihood of two succeeding mutation events increases the rate of successful two-step adaptations.

This effect also emerges for two-step adaptations that do not constitute proper fitness valleys, i.e. for which the intermediate mutant is not maladaptive (Fig 2E; 1.15-fold increase). Under stochastic selection dynamics, even mutants with a slight selective advantage (Fig 2C) have a high chance of going extinct due to random drift. Lineages acquiring sets of synergistic mutations, thus, often do so without prior fixation of each intermediate mutant – via stochastic tunneling^26^. In those regimes, temporal clustering of mutations therefore also increases the speed of adaptation, as confirmed with simulations (Fig S2A-D). Moreover, the proportion of such sets of synergistic mutations acquired via stochastic tunneling rather than sequential fixation also increases with temporal clustering (Fig S2E).

Our results generalize to multi-step adaptations with arbitrarily many steps (SI1). To demonstrate this result, we compared a uniform mutation process with mutation rate *µ* to temporally clustered mutation processes in which mutations emerge exclusively during burst phases that constitute a fraction 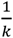 of the time, but at a *k*-fold increased rate, *kµ* (Fig S2A). We showed analytically that in the limit of the average mutation rate *µ* going to zero, the rate *f*_k_ of successful (*n* + 1)-step adaptations is *k*^n^ times higher in the temporally clustered mutation process than in the uniform mutation process. We validated this result with simulations of a Wright-Fisher process for two-step adaptations, *n* = 1, and a range of small values for *µ* (Fig S2F,G), and with numerical calculations of *f*_k_ in a branching process (see SI2-SI3 and Fig S3-S4 for complete characterizations of parameter dependencies). As the mutation rate increases, this fold-increase 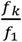 becomes smaller. However, since the absolute rates *f*_*k*_ and *f*_1_ are proportional to *µ*^*n*+1^ (SI1), the absolute effect of temporal clustering on the rate of fixations of mutants with multi-step adaptations increases in *µ* (Fig S2F). Analogously, we found that 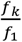 increases with the selective disadvantage that cells in a fitness valley have, while *f*_k_ − *f*_1_ decreases with this selective disadvantage (SI2). In contrast, for low *µ*, and particularly for parameter choices in biologically relevant ranges (*µ* ≤ 10^-7^), the fold-change becomes insensitive to the magnitude of the selective disadvantage so that ever smaller fitness differences suffice to reach high fold-changes, 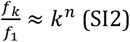. For instance, in a population of constant size that evolves according to a branching process with a birth-death ratio of 1, if intermediate mutations in a multi-step adaptation sequence reduce the birth-death ratio to 0.98, we have 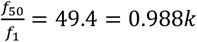 for 2-step adaptations, and 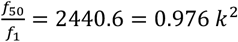 for 3-step adaptations. In light of the large fold-changes in mutation rates during a burst and before or after a burst^10,15,17,18^, our findings suggest that the resulting effect on the dynamics of multi-step adaptation is substantial.

### Temporal clustering in an exploration-exploitation setting

The increased rate of valley crossing is driven by phases of high mutation rates, when the fold-increase in the chance of valley-crossing is larger than the fold-increase in the mutation rate. This effect improves adaptability while the fitness landscape offers scope for multi-step adaptation. However, in fitness landscapes which have global maxima, as the population adapts there is ever less scope for (further) multi-step adaptations, and excessive exploration of the landscape becomes costly.

To investigate the effect of temporal clustering in such an exploration-exploitation setting, we constructed a two-dimensional fitness landscape (Fig 3A) which is randomly re-drawn at regular intervals, mimicking exposure to novel physical environments or drugs during cancer evolution and treatment (Methods). Between subsequent re-drawings, the population may move to a global fitness maximum where any further mutation decreases fitness. We considered mutation processes with different average mutation rates *µ* and different values of the clustering parameter *k* (Fig 3B) and investigated the average fitness of the cell population over long time horizons. As before, the clustering parameter *k* denotes the fold-increase of the mutation rate in burst phases relative to the average mutation rate *µ*. However, rather than fixing the out-of-burst mutation rate at zero, the burst duration is now held constant (Fig 3B) to control for differences in waiting time until a burst occurs.

**Figure 3:**
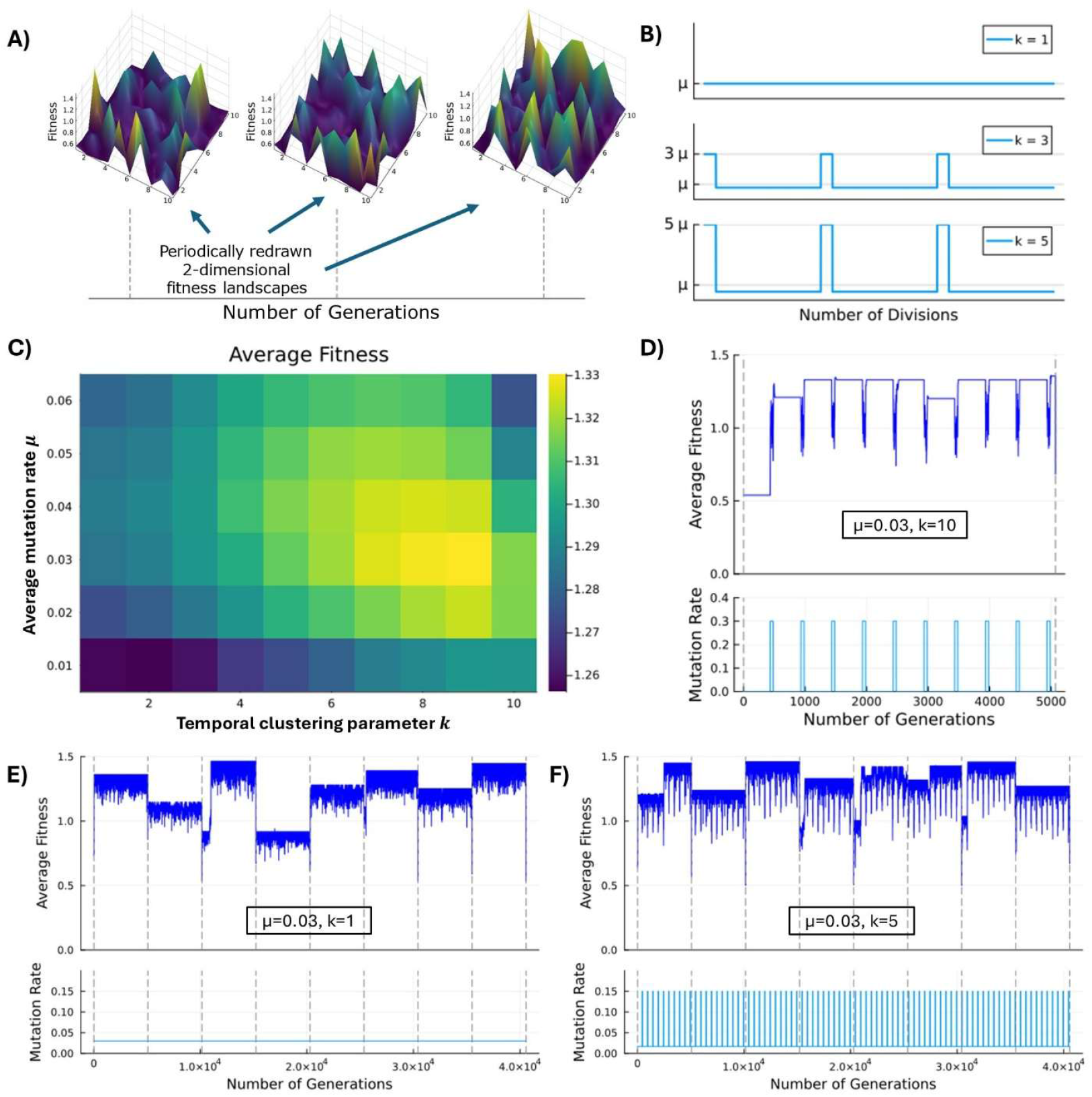
Exploration and exploitation with temporally clustered mutation rates. **(A)** Cells move through two-dimensional fitness landscapes. These fitness landscapes are randomly redrawn every 50714 divisions. **(B)** Mutation rate trajectories are parameterized with a mean mutation rate μ, and a clustering parameter k. **(C)** Average fitness in simulations of a population of 20 cells in a Wright Fisher Process. Simulations were run for 10^8^ division events. **(D)-(F)** Average fitness trajectories for representative snippets of the simulations in **(C)**.

We found that, in this scenario, relative to the uniform mutation process (*k* = 1) increasing *k* initially increases the average fitness of the population for any fixed *µ* (Fig 3C). However, at high *k*, marginal increases of *k* reduce fitness, since high in-burst mutation rates in small populations increase the chance of leaving fitness peaks due to stochastic drift, and low mutation rates outside of bursts delay the acquisition of one-step adaptations. Similarly, for a fixed *k*, increasing *µ* when starting at low *µ*-values increases the average fitness. However, moving further in either of these directions in the parameter space – towards increasing *µ* or towards increasing *k* – eventually brings about a decrease in average fitness reflecting a tradeoff between exploration and exploitation (see SI4 and Fig 3D-F for detailed descriptions and plots of the dynamics). For any fixed *k*, average fitness peaks at intermediate *µ*, and the highest average fitness on the *k*-*µ*-plane is achieved by a *k >* 1. Maximizing average fitness, thus, involves temporal clustering even when there is a trade-off between exploration and exploitation.

### Proxies for valley crossing and for temporal clustering found in patient data

We then sought evidence of the modeling prediction that temporal clustering facilitates multi-step adaptation in patient data, leveraging whole exome sequencing (WES) data from TCGA. We reasoned that among tumors in which similar numbers of mutations had emerged, those tumors with more temporally clustered mutation processes in their evolutionary history would be more likely to have undergone a larger number of multi-step adaptations. Generalizing this idea to enable comparison of tumors which may differ in the number of mutations they have acquired, we reasoned further that tumors with high levels of temporal clustering would have a high fraction of successful multi-step adaptations per attempted multi-step adaptation, i.e. per emergence of a first mutant in a multi-step adaptation sequence. We would, thus, expect analogous ratios of detectable quantities that scale with the numbers of successful or attempted multi-step adaptations to correlate with indicators of fluctuations in the mutation rate history of a tumor such as those found for APOBEC mutagenesis. In particular, we considered the frequency of observing two deactivating mutations in a TSG as an indicator for successful multi-step adaptation, and the total number of observed mutations in a tumor as a quantity that scales with attempted multi-step adaptations.

Before considering the TCGA data, to confirm in silico that such a correlation would indeed emerge under biologically realistic parameters, we performed large-scale simulations of tumor growth from a single cell to up to realistically detectable cell numbers of 10^6^−10^7^ cells^47^ (Methods). We recorded the fraction of mutations that emerged during burst phases, constructed a TSG deactivation score (Fig 4A,B) by counting the number of TSGs (modelled as mutations with synergistic fitness effects) with at least two mutations across the population and divided by the total number of mutations across the genome (Methods). Our simulation results confirmed that these two readouts are correlated (Pearson correlation 0.42, p<0.01; Fig 4C), showing that in silico predictions regarding the likelihood of acquiring 2-step adaptations under mutation processes with more vs. less temporal clustering are well reflected in our score for TSG deactivation.

**Figure 4:**
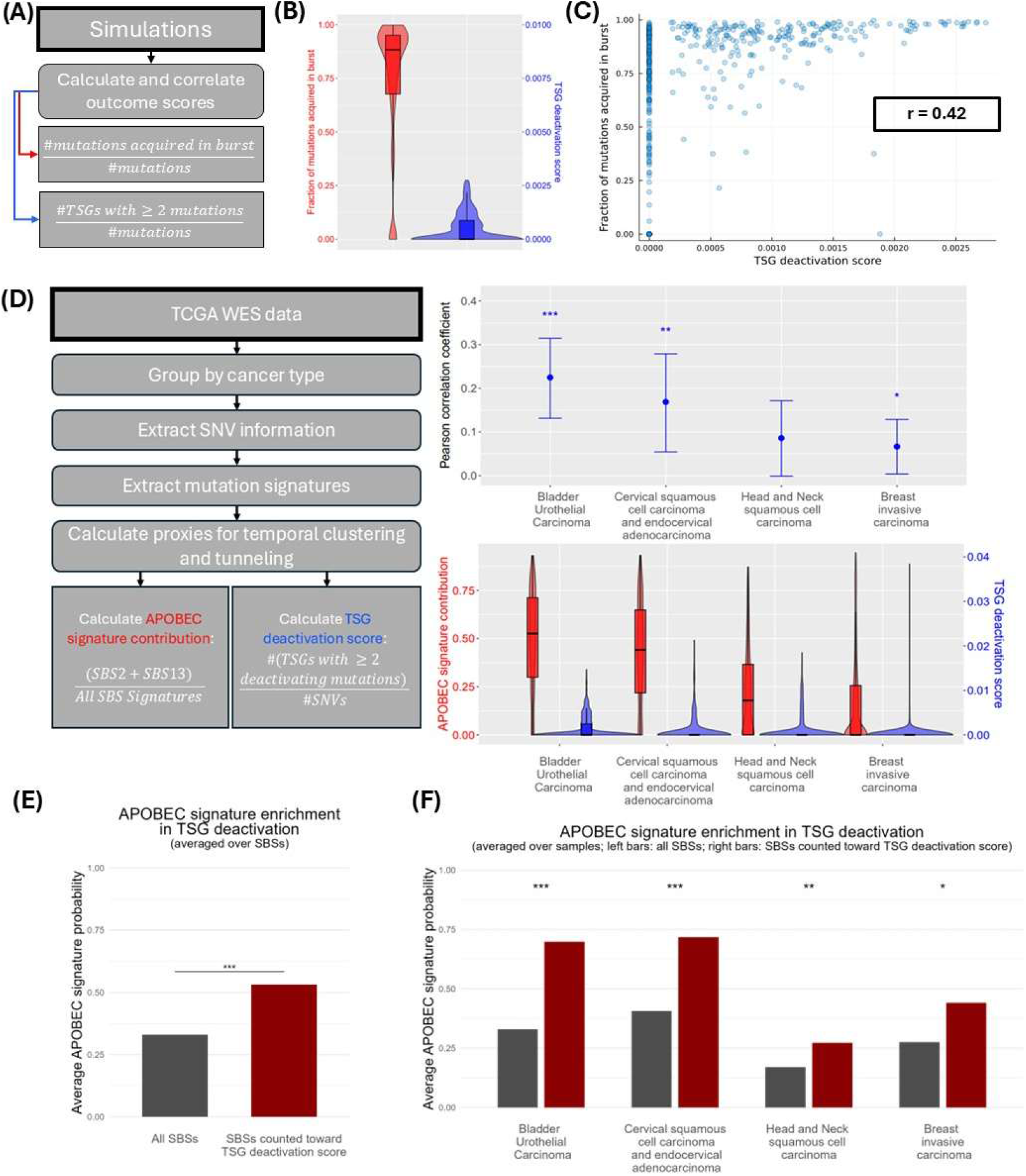
Proxies for valley crossing and temporal clustering in simulations and in TCGA data. **(A)** Sketch of the simulation analysis workflow. **(B)** Distributions of the fraction of mutations acquired during mutation bursts and the TSG deactivation score in simulations. **(C)** Joint distribution of the quantities in (B) indicates strong correlation. **(D)** Schematic of analysis workflow for TCGA data (methods) and results for the four cancer types with highest APOBEC signature contribution. **(E)** Probabilities that single base substitutions (SBSs) in samples of the cancer types in (D) were caused by APOBEC associated signatures SBS2 and SBS13. On the left, these probabilities are averaged over all SBSs. On the right, only those SBSs are considered which are classified as “nonsense” or “missense”, and which appear in a TSG with at least 2 such deactivating SBSs. The average probabilities between both groups are significantly different (t-test, p<0.001). **(F)** Average probabilities analogous to those in panel (D) were constructed for each individual sample and then averaged across samples. Samples without deactivated TSGs were excluded. The average probabilities between groups are significantly different for the four cancer types (^*, **^ and ^***^ respectively indicate that the p-value in a t-test lies below 0.05, 0.01 and 0.001).

Next, to construct similar readouts from the TCGA data, we first performed SBS signature decomposition on each sample and computed the relative contribution of APOBEC-associated mutation signatures (<u>SBS2</u> and <u>SBS13</u>) vs all other mutation signatures^48^ (Methods). Given the evidence for episodic APOBEC mutagenesis ^10,14^, we used this relative contribution as a score for temporal clustering. As a proxy for multi-step adaptations, a list of known TSGs (the TSGene 2.0 database^49^) was considered, and for each TCGA sample, we counted the number of TSGs in this list which harbored at least two single nucleotide substitutions classified as a missense or nonsense mutation (Methods) normalized by the total number of single nucleotide substitutions in that sample. The TCGA WES data did not enable us to determine whether such mutations indeed deactivated different alleles of a TSG, rather than appearing on the same allele. If the likelihood of appearing on the same allele varied with the contribution of the APOBEC signature, this association could confound our results. Since mutations that are interdependently generated on the same copy of a gene would likely appear in close spatial proximity^11,50,51^, we thus verified in a robustness check that our results remain unchanged when filtering out mutations with close-by genomic coordinates (Methods).

We then ranked the different tumor types in TCGA by the average contribution of SBS2 and SBS13 relative to all detected mutation signatures, and for each tumor type, investigated the correlation between our proxies for multi-step adaptations and for temporal clustering. For the top four categories with the highest mean contribution of APOBEC-driven mutagenesis to all mutations, we observed a positive correlation between the two proxies. This correlation was significant at the 5% level (t-test on the Pearson correlation coefficient) for three out of four of these top four categories (Fig 4D, for results for categories with lower APOBEC signature contribution see Fig S5).

Consistent with the hypothesis that this correlation arises because TSG deactivation occurs more readily with APOBEC mutagenesis, we found that mutations which contribute to the TSG-deactivation score on average have a strongly increased likelihood of being caused by APOBEC (Methods, Fig 4E). This finding is consistent across cancer types and independent of whether probabilities are averaged over samples or over individual mutations, suggesting that this result is not driven by outlier samples with many TSG inactivations and many APOBEC-associated mutations (Methods, Fig 4F).

An alternative score for multi-step adaptations, in which the list of TSGs is replaced with a list of pairs of genes with synergistic fitness effects^31^ (Methods) again exhibits a positive correlation with the relative contribution of APOBEC-associated mutation signatures in all categories; this correlation is significant for two of these categories (Fig S5). Additionally, we verified that our results are robust to the choice of correlation metric: using Spearman instead of Pearson correlations yields significant positive correlations across all top four categories discussed above (Fig 4D, S7).

Taken together, our analyses of TGCA data show that tumors with a larger share of APOBEC-associated mutations are enriched for pairs of mutations with synergistic fitness effects. While we cannot exclude the influence of unobserved confounders, given the limitations of the available data, we found that the patterns from our analysis are consistent with the theoretical predictions of our model.

### Stochastic Tunneling vs Clonal Interference during Punctuated Evolution

In tumors evolving at high mutation rates, clonal interference may limit the rate at which a cell population manages to acquire and retain 1-step adaptations (Fig 1F). Previous literature has quantified this limiting effect as a function of the mutation rate and the distribution of fitness values of emerging mutants^52–54^. However, the effects of non-constant mutation rates in this setting have not been elucidated. We thus set out to investigate clonal interference in the setting of temporally clustered mutation rates (Fig 5A).

**Figure 5:**
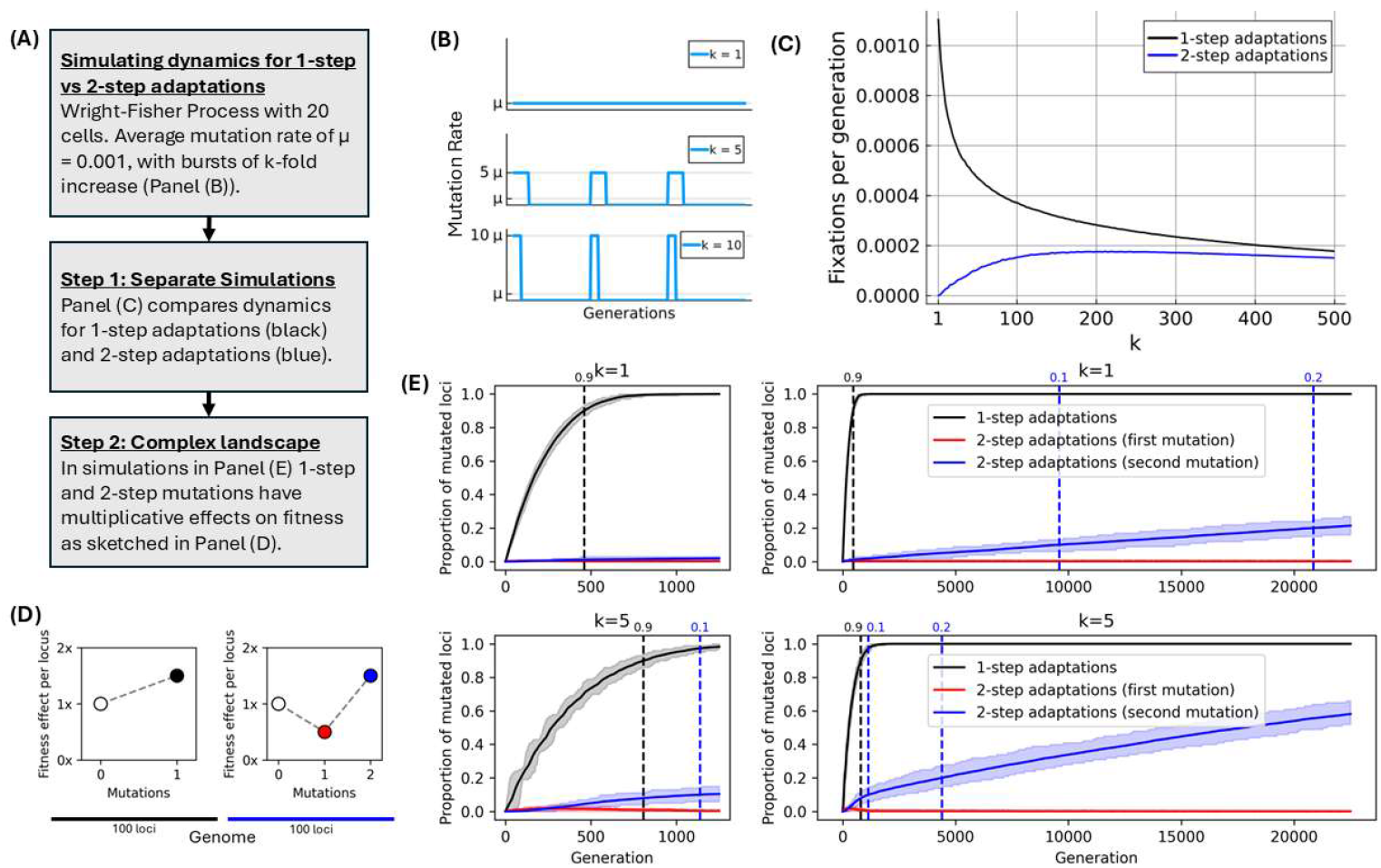
Stochastic tunneling vs clonal interference under temporally clustered mutation rates: **(A)** Schematic of the simulation approach. **(B)** Sketch of the mutation process. **(C)** Simulation results: measuring fixations of higher-fitness mutations per generation as a function of the clustering parameter and of the fitness landscape. In the 1-step adaptation setting fitness is defined as 1.5^#*mutations*^. Fitness in the 2-step adaptation setting is defined as in Fig 2B. Results are averaged over simulation runs with 10^7^ generations. **(D)** Sketch of fitness landscape for simulations shown in (E). Cells start with an unmutated genome of 200 loci. Mutations on each locus have independent multiplicative effects on cell fitness. In the first 100 loci a single mutation confers a multiplicative fitness change of 1.5. In the remaining 100, 2 mutations are required to reach this multiplicative fitness change of 1.5, with the first mutation conferring a multiplicative fitness change of 0.5. **(E)** Simulation results for adaptation with genome sketched in (D). Results are averaged over 100 simulation runs per value of k. Shaded regions indicate 10^th^ and 90^th^ percentiles. Smaller plots (left) are zoomed in versions of the first 1200 generations of the larger plots (right), with identical color-coding.

We first considered a scenario in which cells can only acquire 1-step adaptations and measured the rate at which novel 1-step adaptations reach fixation in the population across simulations with different clustering parameters *k* (Fig 5B). We found that fixation rates quickly decrease as *k* increases (Fig 5C). This pattern emerges because the extent to which clonal interference reduces fixation rates disproportionately increases with the mutation rate; for a fixed average mutation rate, temporal clustering thus increases the effects of clonal interference.

These findings stand in contrast to our results for 2-step adaptations whose rate of fixation increases when temporal clustering is introduced. However, if in-burst mutation rates are sufficiently large such that multiple clones in the population may acquire a 2-step adaptation independently before one of these clones has fixated, effects of clonal interference also play a role for 2-step adaptations. Performing simulations for 2-step adaptations, we found that fixation rates are non-monotone in *k*. While at low *k* increasing *k* leads to a steep increase in the fixation rate, this trend eventually levels off and becomes negative, with further increases in *k* leading to a decrease in the fixation rate (Fig 5C).

Having observed that temporal clustering increases effects of clonal interference but also facilitates stochastic tunneling, we next set out to investigate the relative contribution of these two effects on overall adaptation rates. We considered a setting in which cells can increase their fitness through both 1-step and 2-step adaptations (Fig 5D). We found that 1-step adaptations are acquired more quickly under the uniform mutation process (k=1), while 2-step adaptations are acquired more quickly under a temporally clustered mutation process (k=5; Fig 5E). The relative magnitude of both effects varies with time. As 1-step adaptations spread more readily in the population, differences in the speed at which they fixate manifest early in the dynamics. Over time, cells gradually exhaust the possibilities for 1-step adaptations and the difference between the adaptation rate in both processes becomes dominated by differences in the speed of acquiring two-step adaptations. In this latter phase we observe large differences between the two mutation processes in the expected time until any given proportion of the possible two-step adaptations have spread in the population. Analogous time differences for reaching fixed proportions of one-step adaptations in the initial phase of the dynamics are much smaller (Fig 5E).

In these analyses, parameters such as mutation rates, fitness effects and population sizes were chosen so that simulations remained computationally manageable. Decreasing mutation rates toward more biologically realistic ranges decreases the likelihood that multiple lineages with adaptations emerge simultaneously and therefore weaken the effects of clonal interference. In contrast, larger population sizes and smaller fitness differences increase the likelihood of co-existence between competing lineages as each emerging lineage requires more time to fixate in the population, which strengthens the effects of clonal interference.

We also investigated how this pattern depends on the ruggedness of the fitness landscape (Fig S9A). To this end, we varied the relative proportions of loci with and without fitness valleys and quantified adaptability differences. We found that the duration of the initial phase in which the uniform process yields higher adaptability varies with the ruggedness of the fitness landscape: this phase is roughly three times as long when the proportion of loci with fitness valleys is 5% compared to when it is 95% (Fig S9B-D). However, over this range of proportions, we consistently observed that this initial phase only accounts for a small fraction of the time it takes to acquire all adaptations.

## Discussion

Accumulating evidence suggests that mutagenesis in tumor cell populations proceeds in punctuated bursts rather than gradually; however, the effects of such punctuation on the ability of a tumor cell population to acquire and retain fitness-enhancing adaptations remains incompletely understood. Here we set out to investigate the evolutionary dynamics of these processes using mathematical modeling and genomics data analysis of human tumors.

We found that when a population acquires multi-step adaptations via stochastic tunneling, the rate of adaptation is substantially enhanced by punctuation. Stochastic tunneling is the predominant mode of evolution in large populations that traverse fitness valleys. However, stochastic tunneling also occurs when the necessary mutation steps to reach a substantial fitness advantage are not individually maladaptive, i.e. if steps do not constitute a fitness valley but confer a slight fitness advantage. Punctuated evolution therefore facilitates the accumulation of sets of mutations that jointly and synergistically confer a fitness advantage, while the fitness effect of carrying only a subset of these mutations can range from being strongly maladaptive to being moderately adaptive. For the limit of low average mutation rates we show analytically that the rate of stochastic tunneling under a temporally clustered process is *k*^*n*^ times larger than under the time-invariant process, where *k* is the fold-increase in the mutation rate relative to the average mutation rate during mutation bursts in the temporally clustered process, and *n* is the number of mutation steps that is required to exit the fitness valley. Moving towards higher average mutation rates, this fold-change in the stochastic tunneling rates decreases, but the absolute difference between the rates increases. Investigating these relative and absolute effects in a branching process as a function of the birth-death ratio elucidated a similar pattern: While the fold-change is highest if lineages in a fitness valley have a low birth-death ratio, we observed the largest difference between rates of stochastic tunneling at birth-death ratios close to one.

Applied to cancer evolution where cells’ birth-death ratios are influenced by a variety of internal and external factors, these findings highlight that for any given multi-step adaptation sequence, the magnitude of the facilitating effects described here likely varies depending on whether the tumor is in a phase of constant or growing population size.

While temporal clustering facilitates multi-step adaptations, it also impedes 1-step adaptations due to clonal interference. We examined the interplay between these contrasting effects in a setting in which cells can acquire both types of adaptations. We showed that the relative importance of both effects varies over time. Since 1-step adaptations tend to be acquired more readily, clonal interference matters most in the early phase of the process. As the share of possible 2-step adaptations relative to 1-step adaptations increases, differences in the tunneling rate become the more relevant determinant for the speed of adaptation. A uniform mutation rate, thus, makes a population more efficient at finding local fitness maxima. However, temporal clustering allows the population to more quickly move between local maxima, thus speeding up the search for a global maximum.

While our theoretical results are easily verified in simulation settings where we can choose a fitness landscape, applying these results to real data remains challenging because of the inherent complexities in determining fitness effects in biological systems. Estimating fitness effects of individual mutations requires either intricate experimental approaches^55^ or large patient cohorts to control for differences in the (epi-)genetic background against which these mutations emerge; such estimations become considerably harder when considering joint fitness effects of sets of multiple mutations.

To circumvent these challenges, we focused on TSGs as a representative set of gene pairs with synergistic effects and on one mutation process known to tend to fluctuate over time – APOBEC-driven mutagenesis. Moreover, we restricted attention to single-nucleotide substitutions as the principal mode of APOBEC-driven mutagenesis and the context in which mutation signatures are best characterized, and we did not consider other classes of mutations such as copy number changes or epigenomic modifications. Consequently, we observed only a subset of groups of mutations with synergistic fitness effects in the data and possibly only a small fraction of the variability in punctuation between different samples. Nevertheless, we found that these scores significantly correlate, which aligns with our model predictions. However, given this narrow scope, the generalization of our model towards a full quantitative estimate of the relevance of punctuation to adaptation dynamics in cancer requires additional investigations.

The observed empirical patterns may in part reflect the influence of unmeasured confounders. Rates of multi-step adaptation are determined not only by temporal clustering, but also by factors such as average mutation rates, fitness landscapes, and population-size trajectories. Structural differences in these factors between tumors with high or low APOBEC signature contributions therefore represent potential sources of confounding (for a review of known APOBEC-related biological mechanisms and clinical associations, see^60^). Additionally, we cannot measure rates of multi-step adaptation directly, and therefore need to consider a proxy; to this end, we chose a score that relates deactivating TSG mutations to the absolute mutation count. Structural differences affecting the relative rates of deactivating TSG mutations, such as APOBEC-related enrichment for nonsense mutations^59^, compared to other mutations can therefore also influence our empirical findings.

Our results have implications for both mechanistic and statistical modeling of tumor evolution. A key quantity in mechanistic models is the rate at which different types of adaptations arise; our results show that this rate depends on the temporal dynamics of the mutation process. Accounting for this effect might be particularly important when modelling the emergence of treatment resistance in models used to optimize treatment schedules^56^. There is evidence linking APOBEC mutagenesis to resistance in breast cancer^61^. Our results suggest that APOBEC, and other punctuated mutation processes, are likely to be relevant more generally to settings in which developing resistance requires multi-step adaptation. Analogously, in statistical models, incorporating measures of punctuated mutagenesis may improve the ability to predict treatment outcomes by accounting for how these temporal dynamics shape adaptability.

## Methods

### Simulations of a Wright-Fisher and a branching process in an unbounded fitness landscape

In the simulations presented in Fig 2 and Fig S1, mutations occur between selection steps and are independent of divisions. In the Wright-Fisher process simulations (Fig 2 C) with a temporally clustered mutation rate, the out-of-burst mutation probability per cell and per update step (between two divisions) was set to 0.01 and was increased to 0.1 in bursts. For the corresponding uniform mutation rate trajectory, this probability was simply set equal to the total amount of mutations that occurred in the simulations under the temporally clustered mutation rate, divided by the total number of update steps.

Mutation rates in the branching model simulations (Fig S1) were chosen analogously. However, to achieve temporally equidistant mutation bursts in the branching process simulations, we scaled the burst duration and the time between bursts by the population size.

### Simulations Wright-Fisher Process in exploration/exploitation setting

To produce the results shown in Fig 3, we performed agent-based stochastic simulations of a Wright-Fisher Process with 20 cells. Cells were characterized by their location on a discrete fitness landscape (a 30 by 30 two-dimensional lattice). The fitness values *f* associated with the positions on the lattice were drawn independently at random as *f* = 0.5 + *x*^4^ where *x* ∼ *U*[0,1] is distributed uniformly between zero and one. The lower bound of 0.5 ensures that cells at all positions have a non-negligible probability of dividing.

Raising *x* to the fourth power creates a landscape with few peak-positions surrounded by many valley-positions with little variation in fitness.

In each selection step, one cell was randomly chosen to divide with probability proportional to its fitness, and replaced a cell which was drawn uniformly at random. Mutations occurred between consecutive selection steps, and the mutating cell was chosen uniformly at random amongst all cells in the population (independently of the preceding selection step). If a cell mutated, it would move to a location in the fitness landscape chosen uniformly at random from the set of (at most 8) locations in the Moore neighborhood of its current location.

The rate at which mutation events occurred was governed by the parameters *µ* and *k*. Mutation bursts lasted 10^2^ division events and started every 10^3^ division events. Fitness landscapes were redrawn every 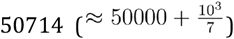 division events, to periodically vary the relative timing of mutation bursts and re-drawings of the fitness landscape. In this manner, we avoid artifacts in our results caused by phase-alignment of bursts and re-drawings.

### Larger-scale simulations

We simulated tumor evolution as a branching process, starting from a single unmutated cell, up to a randomly drawn target cell number between 10^6^ and 10^7^ cells. We assume a constant death rate, so that over the course of a simulation the probability that the next event is a death event rather than a division event remains fixed (at a value of 0.35). In case of a death event, one cell picked uniformly at random is removed from the population. In case of a division event, one cell is chosen to divide with probabilities proportional to its fitness.

We assume that the number of mutations per division follows a binomial distribution *Bin*(10^5^,*µ*), and for each simulation we randomly draw a baseline mutation probability *µ* uniformly between 0 and 3·10^−5^. In mutation bursts, this probability gets multiplied by a factor of 20 or 50, again chosen randomly for each simulation. The population enters a burst phase with probability 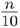 per division, where *n* is the current population size, and exits a burst phase with probability 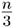, so that in expectation bursts phases last 3 generations, and start every 10 generations. The first cell in a simulation is initialized to be in a burst with probability 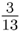 consistent with the expected time spent in bursts given those parameters.

The fitness effect of mutations is modeled as follows. The starting cell has a fitness of one. Each new mutation has an additive effect on the cell’s fitness. We assume that mutations occur uniformly at random across the genome and that multi-step adaptations can occur in a fraction *θ* = 0.01 of the genome, roughly aligning with the ratio of the number of TSGs vs the total number of genes in the human genome. We subdivide this part of the genome into 200 TSGs which are all hit by a mutation with equal probability. For each individual TSG, the first mutation reduces a cell’s fitness by 0.05. The second mutation increases fitness by 0.15, and all further mutations are fitness neutral. For the remaining (1 − *θ*) fraction of the genome, we assume that mutations are fitness-neutral with a probability of 0.9, and that fitness effects are otherwise drawn from a Gaussian with mean equal to −0.005 and standard deviation equal to 0.005.

To produce the results in Fig 4A-B, we keep track of all mutations that arise in a population, and remove all mutations with less than 1% variant allele frequency from the output, as those would be unlikely to be detected in WES. Moreover, we keep track of the fraction of mutations that a sample acquired during a mutation burst. For each simulation, we then count the number of TSGs for which there are at least two mutations found in the population and divide this by the total number of mutations in the sample.

### Analysis of TCGA WES data

Whole-exome sequencing data was acquired from The Cancer Genome Atlas. We performed mutation signature decomposition using the *cosmic fit()* function in python from the package SigProfilerAssignment^57^ version 0.1.8 with cosmic version 3.4. Moreover, we used the mutation-level signature probabilities generated via SigProfilerAssignment to compute mean probabilities of signatures SBS2 and SBS13 for individual SBSs (Fig 4 D,E).

To compute our TSG deactivation score, we filtered the SNV data for missense and nonsense mutations in genes belonging to the TSGene 2.0 database^49^. For each sample we calculated the number of TSGs with at least two mutations and divided by the total number of SNVs.

Analogously, to construct our synergistic mutations score, we filtered for missense and nonsense mutations in frequently co-mutated gene pairs identified by Gu et al.^31^, and for each sample divided the number of pairs in this list with mutations in both genes by the number of SNVs. ^29^This enrichment for co-occurrences of mutations in both genes in a pair suggests that mutations in the two genes tend to have a synergistic fitness effect. For each sample in the TCGA WES data we thus computed the number of gene-pairs from this list for which there is at least one non-synonymous single nucleotide substitution mutation in each of the two genes, and divided this number by the total number of single nucleotide substitutions in the sample.

As a robustness-check, we investigated whether the results change if we require a minimum distance between the genomic locations of the mutations in TSGs that we count to our TSG deactivation score. APOBEC has been linked to clustered mutagenesis through mutation processes of kataegis and omikli^11,50,51^. If samples with higher APOBEC activity have higher rates of having multiple close-by mutations on the same allele, and if such a sample has multiple deactivating mutations in a TSG, these inactivating mutations might be more likely to be localized on the same allele compared to samples with lower APOBEC activity. Since we use appearances of more than one inactivating mutation as a proxy for bi-allelic deactivation of TSGs, such a pattern would bias our results. To account for that, we explored filtering TSG mutations in the data based different minimum-distance-thresholds ranging from 1 to 100 bp and found no impact on our results. We did not perform any analyses with allele-specific mutation information, as determining allele-specific TSG inactivation relies on informative co-occurring mutations. For any given mutation, tumors with lower mutation burden are less likely to harbor such co-occurring mutations and preferentially filtering out observations in tumors with lower mutation burden would have biased our findings.

## Supplementary Information

### SI1 Derivations of results on multi-step adaptation rates in the limit of infrequent mutations

In the main text, we discussed differences in the rate of multi-step adaptations *f*_*k*_ as a function of a temporal clustering parameter *k*. Here we formalize this discussion and derive the result presented in the main text. We consider processes in which multi-step adaptations occur via stochastic tunneling rather than sequential fixation, i.e. processes in which mutants with a selective disadvantage are negligibly unlikely to reach fixation. We assume that mutations happen sufficiently infrequently, so that we can neglect valley crossing scenarios in which the same mutation in a multi-step adaptation sequence occurs multiple times. Additionally, we require that valley crossing happens solely via stochastic tunneling, i.e. that the probability that a maladaptive mutant fixates is vanishingly low.

We denote the average rate at which cells acquire mutations by *µ*, and compare dynamics under different mutation processes which we index by a temporal clustering parameter *k* ≥ 1. Starting every *d* units of time, a process with parameter *k* undergoes a mutation burst lasting 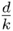 units of time, during which mutations occur at a rate *kµ*. Outside of bursts, the mutation rate is 0. We assume that the time 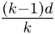 between subsequent burst phases is sufficiently long such that we can restrict attention to multi-step adaptations that occur during a single burst, and that the duration of each burst 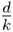 is sufficiently long relative to the time that it takes to cross a fitness valley, so that valley crossings, which fail because the burst phase ended but which would have been successful, have a negligible effect on *f*_*k*_.

As stated in the main text, with this setup we can show that for any population dynamics process for which the above limits can be motivated, the fold-increase in the rate *f*_*k*_ at which (*n* + 1)-step adaptations occur in a process with clustering parameter *k* relative to the uniform process approaches *k*^*n*^.

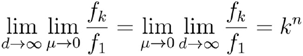

Deriving this result for any selection process indeed becomes straightforward, if it can be motivated that the mutation rate does not affect the fate of any mutant lineage once the first mutant in this lineage has emerged.

Somewhat more formally, we index the steps in an n-step adaptation sequence by *I* ∈ {1,…,*n*}, and denote the expected size of the lineage descending from mutant *i* by *Y*_*i*_(*t*). This quantity *Y*_*i*_(*t*) of course depends on the specificities of the selection process, but for our derivations it suffices to only require that *Y*_*i*_(*t*) does not depend on the mutation rate.

Since all intermediate mutants have a selective disadvantage, and since we assume that we are in a parameter regime in which the likelihood that a disadvantageous mutant fixates is negligibly low, it follows that the integral 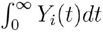 converges.

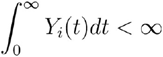

In the limit of a low (and for now time-invariant) mutation rate *µ*, the probability that the *i*’th mutant produces a further mutant *P*(*i* → *i* + 1) is simply the product of the mutation rate and this integral.

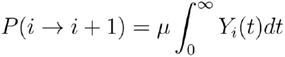

The probability that the first mutant spawns a sequence of *n*+1 mutants can, thus, be written as follows.

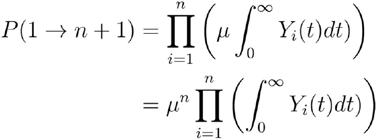

Lastly, we use 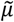 to denote the rate at which new mutants with only one mutation emerge. For a given selection process, this rate may vary with the population size. For the Wright Fisher models of constant population size *N*, this expression simplifies to 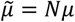. We arrive at the following formulation for the rate at which new advantageous mutants emerge *f*_1_.

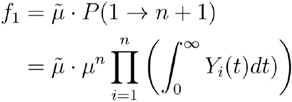

Finally, we can introduce our clustering parameter *k* to scale both *µ* and *µ*-, and multiply by a factor of 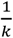 to arrive at the average rate of valley-crossing in the temporally clustered process with parameter *k*.

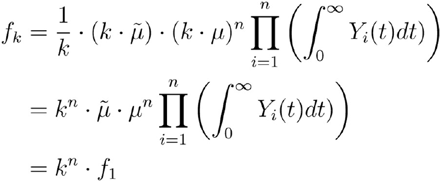

One consequence of our assumption that none of the maladaptive intermediate mutants reaches fixation is that the composition of the population once the final mutant emerges in the limit of *µ* → 0 does not depend on the mutation rate. For population dynamics models with a constant population size in which the probability that an emerging mutant (absent further mutation events) reaches fixation only depends on this composition, such as the Wright-Fisher process, we can interpret 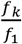 therefore also as the ratio of the rates at which mutants with *n*+1mutations fixate.

Analogously, in certain models of branching evolution in which the prospects of one branch do not depend on other co-evolving branches, such as the Galton-Watson process^58^, whether the lineage of an emerging mutant with (*n* + 1) mutations survives is independent of the mutation rate in the limit of *µ* → 0. The ratio 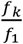 therefore also reflects the ratio of the rates at which surviving lineages with (*n*+1) adaptations arise in such models. We provide thorough treatment of dynamics in such processes below (SI2).

The above result suggests that the effect of temporal clustering on relative rates of valley crossing is substantial, and gets exponentially stronger the wider the fitness valley is (*n*). We validated this result with simulations of a Wright-Fisher process for a range of small values for *µ* (Fig S2 B-E). As we move away from the limit of rare mutations, the relative effect of temporal clustering 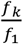 gets smaller, as will be discussed below. However, as the absolute rates *f*_*k*_ and *f*_1_ are proportional to *µ*^*n*+1^, the absolute effect of temporal clustering on the rate of fixations of mutants with multi-step adaptations initially increases in *µ* (Fig S2 D).

### SI2 Dynamics in a Branching Process

In the Galton-Watson process, a branching process with constant birth and death rates, it is possible to derive explicit formulas for the probability that a lineage will acquire a mutation before going extinct (often termed ‘evolutionary rescue’^62^). Formulating this probability for the lineage of the first mutant in a 2-step adaptation sequence thus allows us to explore numerically how the magnitude of the effects that we describe behaves across a wide range of parameter combinations, which is especially attractive for parameter regions for which simulations become prohibitively expensive.

We thus investigated this probability of evolutionary rescue, which translates to *f*_k_/*µ* in our model, as a function of *k, µ* and the birth/death ratio (Fig S3A-C). Note that the birth/death ratio that we considered here is that of the first mutant, not that of the wild type cell. Unsurprisingly, we found that as *µ* decreases, rescue probabilities for any *k* converge to zero for subcritical processes (birth/death ratios < 1). For supercritical processes (birth/death ratios > 1), rescue probabilities converge to one minus the extinction probability of the lineage – rescue only happens if the process expands indefinitely. Throughout, rescue probabilities of course increase in the expected time that the lineage persists, i.e. they increase with the birth death ratio. However, this increase is most steep for large *µ* and flattens for the subcritical range as *µ* decreases.

For the ratio of these escape probabilities for different *k* relative to that of the uniform process, i.e. for 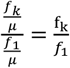, we found that in the subcritical regime, as *µ* decrases 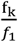 approaches *k*, consistent with our derivations in the previous section (SI1). In the supercritical regime, the condition 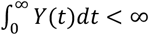 used in our derivations above is violated – lineages have a non-zero chance of branching out indefinitely. Here we saw that as *µ* decreases towards zero, 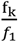 approaches 1 (Fig S3D-F).

Moreover, we found that 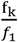 consistently decreases with increasing birth-death ratio. As a result of the interplay between this decrease in 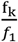 and increase in the absolute rescue probability, we found that the absolute effect, *f*_k_ − *f*_1_, is highest for birth/death ratios close to one (Fig S3G). These results remain as *µ* decreases (Fig S3H-I) but the magnitude of the absolute effect of course shrinks, as rescue of lineages that would otherwise go extinct (all subcritical lineages, and those lineages in the supercritical regime that go extinct by chance) generally becomes unlikely at small values of *µ*.

### SI3 Dynamics as a function of time between bursts

In our derivations, we assumed that bursts are sufficiently spaced out in time, and last sufficiently long so that we can restrict attention to mutant lineages that emerge and go extinct within the duration of a single burst (*d* → ∞). To investigate how effect sizes behave away from this limit, as a function of the time between bursts, we numerically approximated rescue probabilities for finite *d*.

Specifically, we approximated the Galton Watson process by a Markov process with finitely many states *s* ∈ {0,1, …, *N, rescue*}, starting with *s*(0) = 1, where *s* ∈ {0,1, …, *N*} corresponds to the number of viable cells. We considered 0, *N* and *rescue* as absorbing states, and otherwise modelled transitions to states *s* − 1 or *s* + 1 as occurring at rates *s* ⋅ *r_d_* and *s* ⋅ *r_b_* respectively with constant birth and death rates *r*_d_ and *r_b_*, and transitions to the *rescue* state as occurring at rate *s* ⋅ *k* ⋅ *µ* within bursts and at rate zero otherwise. We iteratively solved the corresponding forward equations of this continuous time Markov process numerically for subsequent cycles of in-burst and out-of-burst periods, until the summed mass on states {1, …, *N* − 1} fell below a tolerance threshold, and we considered the joint mass on states *N* and *rescue* as approximation for the rescue probability. For our results in Fig S4 we used N=300 and a tolerance threshold of 10^-10^.

Generally, we showed that differences in multi-step adaptation rates emerge because lineages with a first mutation experience higher mutation rates under a temporally clustered mutation process compared to a uniform mutation process (SI1). As the burst duration and the time-interval between subsequent bursts decrease, this expected difference in mutation rates for mutant lineages diminishes because of the increased likelihood that a mutant lineage that emerged in a burst also experiences a non-burst phase. Consequently, we found that lowering *d* diminishes the effect of temporal clustering on the probability of producing a second mutant (Fig S4 A-C). This pattern manifests both in the relative effects, *f_k_*/*f*_1_ (Fig S4 D-F) and in the absolute effects *f_k_* − *f*_1_ (Fig S4 H-J). However, for the absolute differences *f_k_* − *f*_1_, we found that the strength of this dependency varies non-monotonously with the birth-death ratio. While absolute differences under large *d* peak close to a birth-death ratio of one (Fig S3, Fig S4J), we also observed the strongest decrease in absolute differences at these values for the birth-death ratio, so that under low *d* this peak turns into a local minimum of the absolute-difference curve (Fig S4H,I). This effect arises because the expected lifetime of a mutant lineage before going extinct increases steeply when moving from a subcritical process (birth-death ratios < 1) to a critical process (birth-death ratio = 1), making such lineages with higher birth-death ratios more sensitive to changes in mutation rates that occur long after the lineage first emerged.

### SI4 Dynamics in exploration-exploitation setting

Our simulations show that for given *µ* the effect of increasing the clustering parameter *k* on the average fitness becomes negative at some *k* (Fig 3C). In the simulations in this section, this effect is driven by two main factors. Firstly, after a re-drawing, the population often is not in a local maximum of the fitness landscape, and hence may be able to increase its fitness by single mutations (1-step adaptations), without tunneling. Waiting for a burst to occur and thereby delaying such local explorations decreases average fitness. Second, once the population has found a peak in the fitness landscape, having very high mutation rates in bursts temporarily scatters cells to lower points in the landscape, and may even cause drift to points of lower fitness. Both of these effects become apparent when considering a representative snapshot of the simulations (Fig 3D). The misalignment of re-drawing and burst causes the population to remain in a valley for several divisions until the first burst occurs. Moreover, relative to similar snapshots of simulations with lower *k* (Fig 3E,F), once the population has reached a peak, the scattering during bursts at higher *k* leads to sharper decreases in fitness, and the population may even remain in a lower fitness point after a burst (Fig 3D).

This increased scattering can also be seen when going from *k* = 1 to *k* = 5 (Fig 3E,F). However, at *k* = 1, the population spends much time in local maxima, as large jumps in fitness only occur right after re-drawings, whereas at *k* = 5 jumps occur also long after the landscape was re-drawn and the population found a local maximum. These later jumps to higher points in the landscape correspond to tunneling events.

### SI5 TSG-deactivation and ROS-associated mutagenesis

Following a reviewer suggestion, we also explored the contribution of mutation signatures attributed to reactive oxygen species (ROS; Fig S8). However, we found that these signatures account for only small fractions of the total observed mutation signatures, suggesting that these mutagenic processes would not account for much of the fluctuation behavior of overall mutation rates and we saw no consistent correlations with TSG inactivation.

**Supplementary Figure 1:**
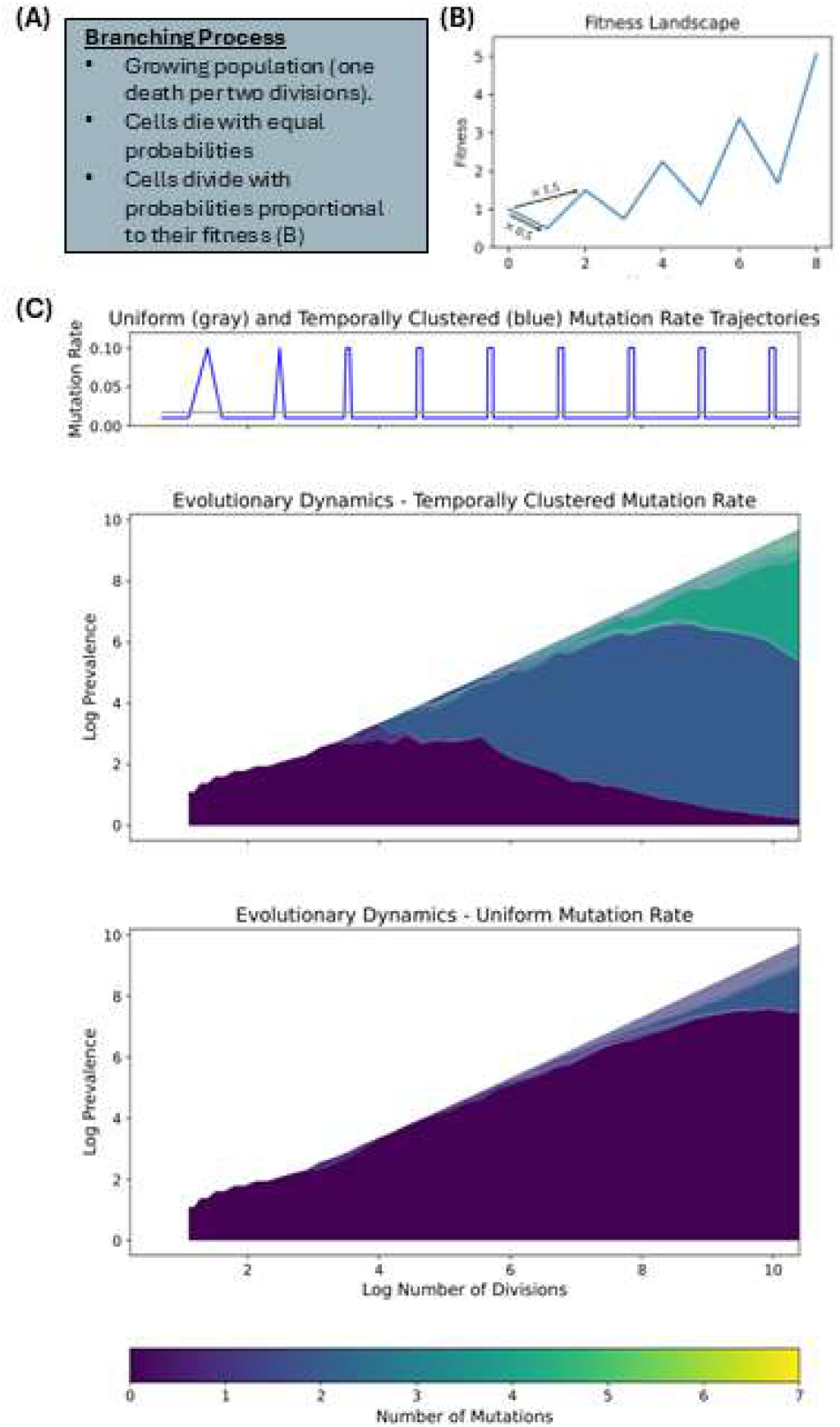
Simulation results: valley-crossing under uniform vs. temporally clustered mutation rates in branching process. **(A)** Schematic of fitness landscape as in Fig 2. **(B)** Schematic of Branching Process model. **(C)** Simulation results for the Branching Process model. The mutation rate trajectories in (C) are chosen such that the total expected number of mutations under the uniform trajectory is identical to that under the temporally clustered trajectory.

**Supplementary Figure 2:**
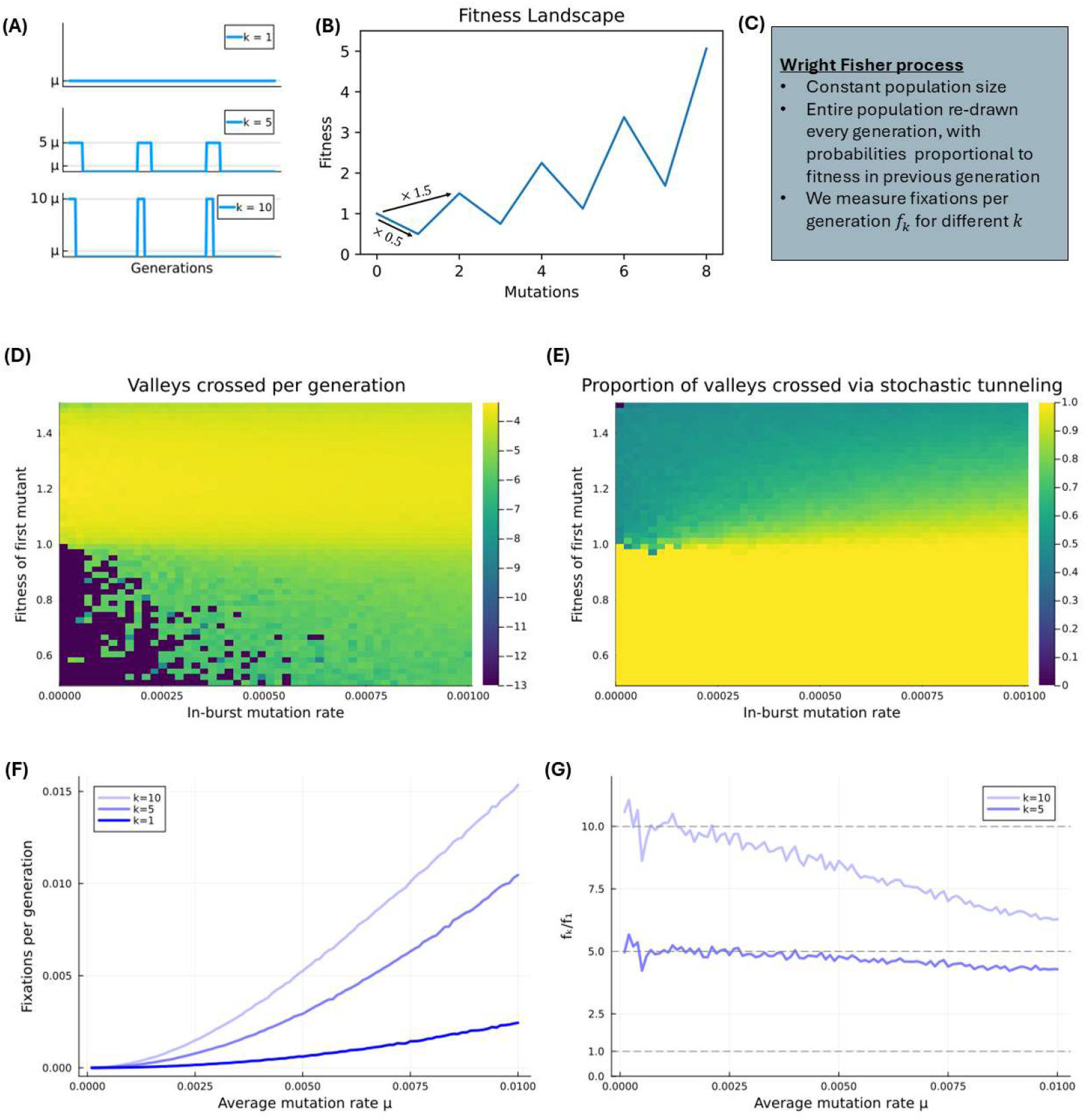
Effect of temporal clustering on stochastic tunneling rates as a function of the mutation rate. **(A)** Sketch of the mutation process with clustering parameter k. **(B)** Sketch of the fitness landscape as in Fig 2. **(C)** Description of the simulations. **(D)-(E)** Simulation results of a Wright-Fisher process with 100 cells over 10^6^ generations with a fixed average mutation rate of10^-5^ per cell per generation and varying temporal clustering k. The fitness is as in (B), with subsequent peaks difering in fitness by a factor of 1.5. However, on the y-axis in (F) and (G) we vary the fitness of the valleys relative to the preceding peak from 0.5 (as in (B)) to 1.5. Panel (D) shows the number of valleys crossed per generation (color bar in log10 values), where contrary to the previous panels it is no longer true that every valley crossing leads to a fixation – the next mutant might emerge before that. Panel (E) the share of these valley crossings that occur without fixation of the first mutant.**(F)** Simulation results of the process described in (A)-(C). Each simulation ran for10^5^/(*kµ*) generations. The y-axis measures fixations of adaptive mutations per generation (see Fig 5D). **(G)** Fixations per generation relative to the uniform mutation process (k=1) for k=5 and k=10.

**Supplementary Figure 3:**
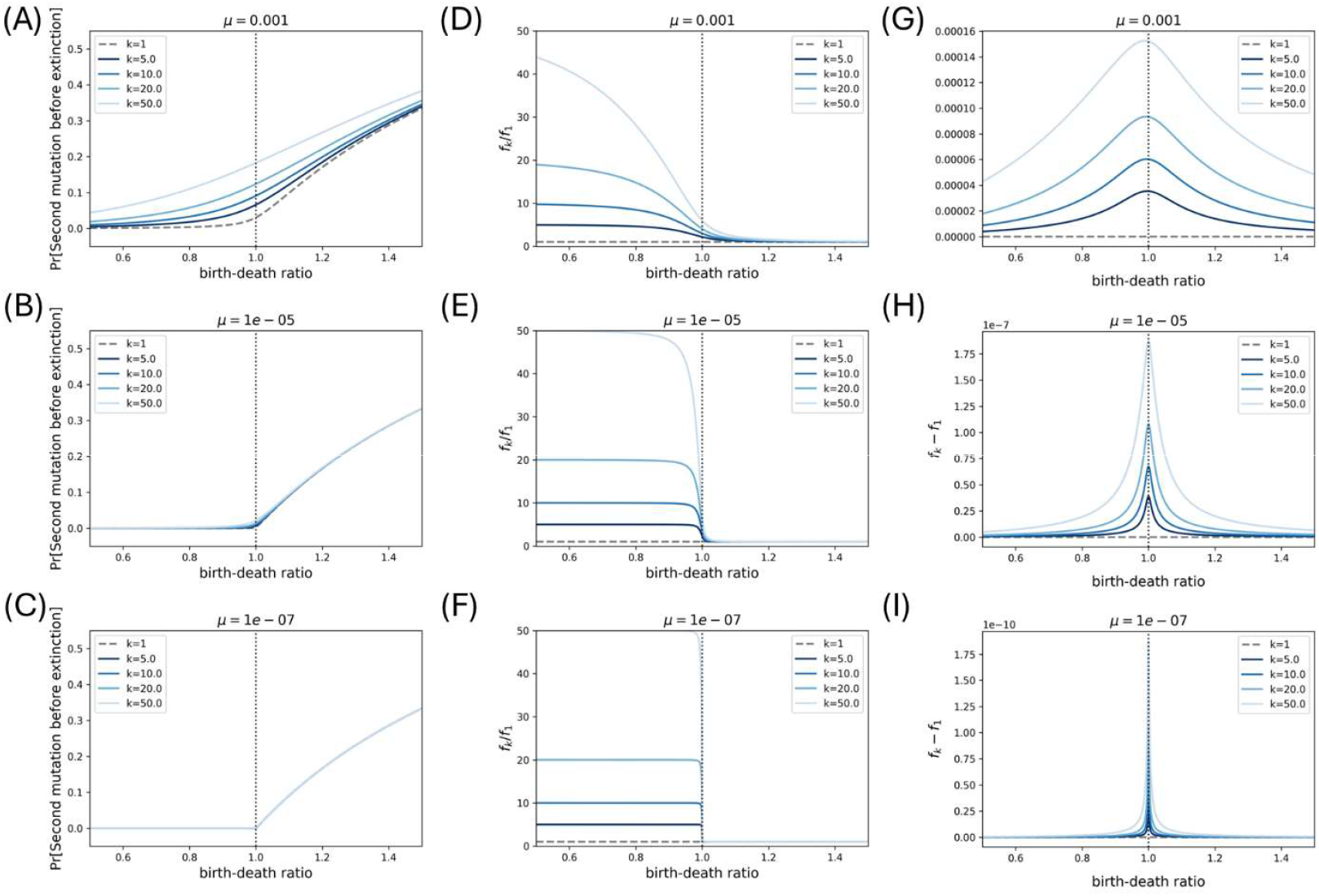
Effect sizes for 2-step adaptations in a Galton-Watson process as a function of the mutation rate and the birth-death ratio. Each row corresponds to a unique value for *µ*. **(A)-(C)** Probabilities that a mutant lineage will give rise to a second mutant. **(D)-(F)** Ratios, *f_k_*/*f*_l_, of the rates at which lineages with one mutation produce a lineage with both mutations for processes with different clustering parameters *k*. **(G)-(I)** Differences, *f_k_* − *f*_l_, of the rates at which lineages with one mutation produce a lineage with both mutations for processes with different clustering parameters *k*.

**Supplementary Figure 4:**
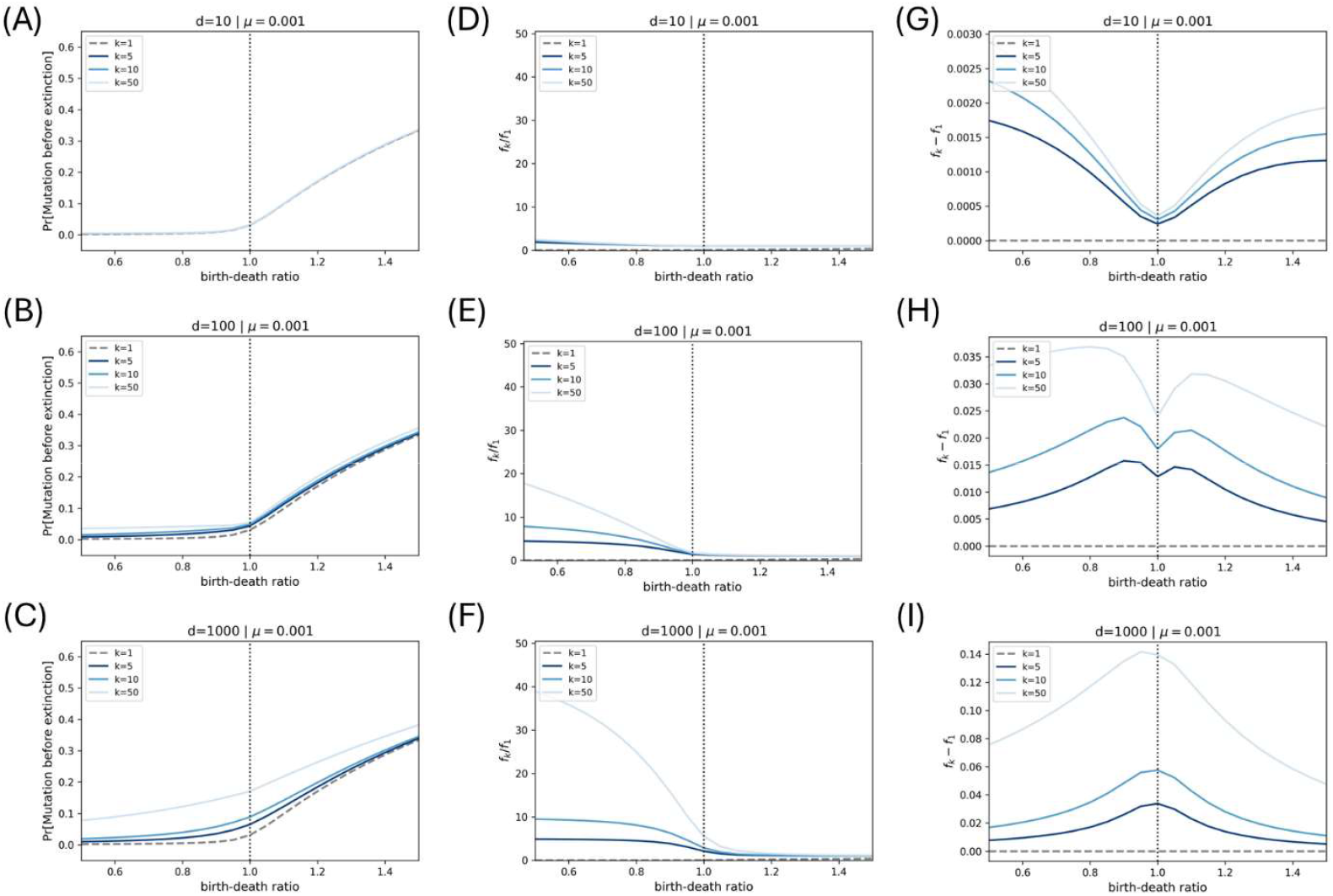
Effect sizes for 2-step adaptations in a Galton-Watson process as a function of the distance *d* between subsequent burst phases. Each row corresponds to a unique value for *d*. **(A)-(C)** Probabilities that a mutant lineage will give rise to a second mutant. **(D)-(F)** Ratios, *f_k_*/*f*_l_, of the rates at which lineages with one mutation produce a lineage with both mutations for processes with different clustering parameters *k*. **(G)-(I)** Differences, *f_k_* − *f*_l_, of the rates at which lineages with one mutation produce a lineage with both mutations for processes with different clustering parameters *k*.

**Supplementary Figure 5:**
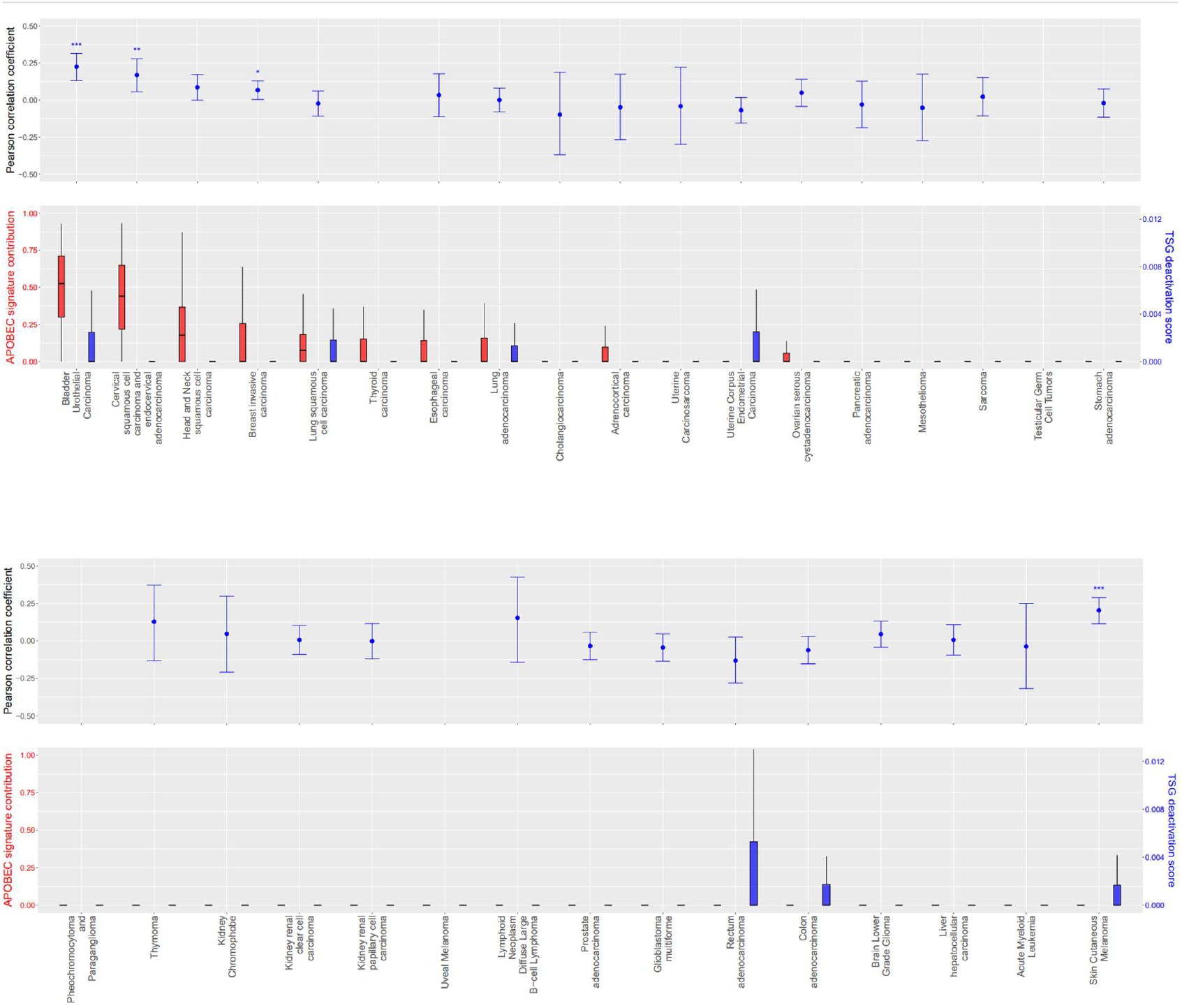
TSG deactivation scores in TCGA. Distribution and correlation of APOBEC signature contribution and TSG deactivation scores as in Fig 4, sorted by cancer type.

**Supplementary Figure 6:**
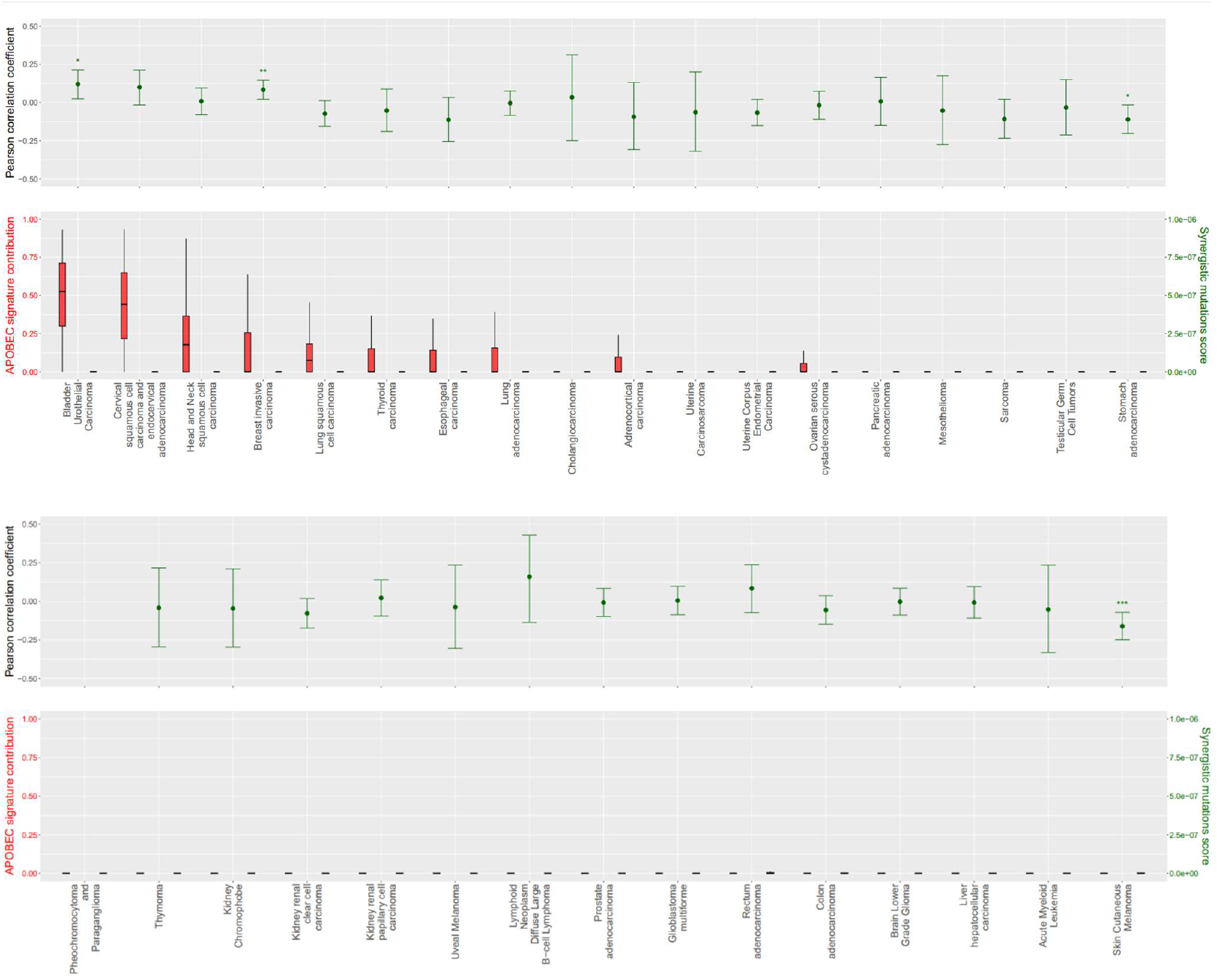
Synergistic mutation scores in TCGA. Distribution and correlation of APOBEC signature contribution and synergistic mutation scores, sorted by cancer type.

**Supplementary Figure 7:**
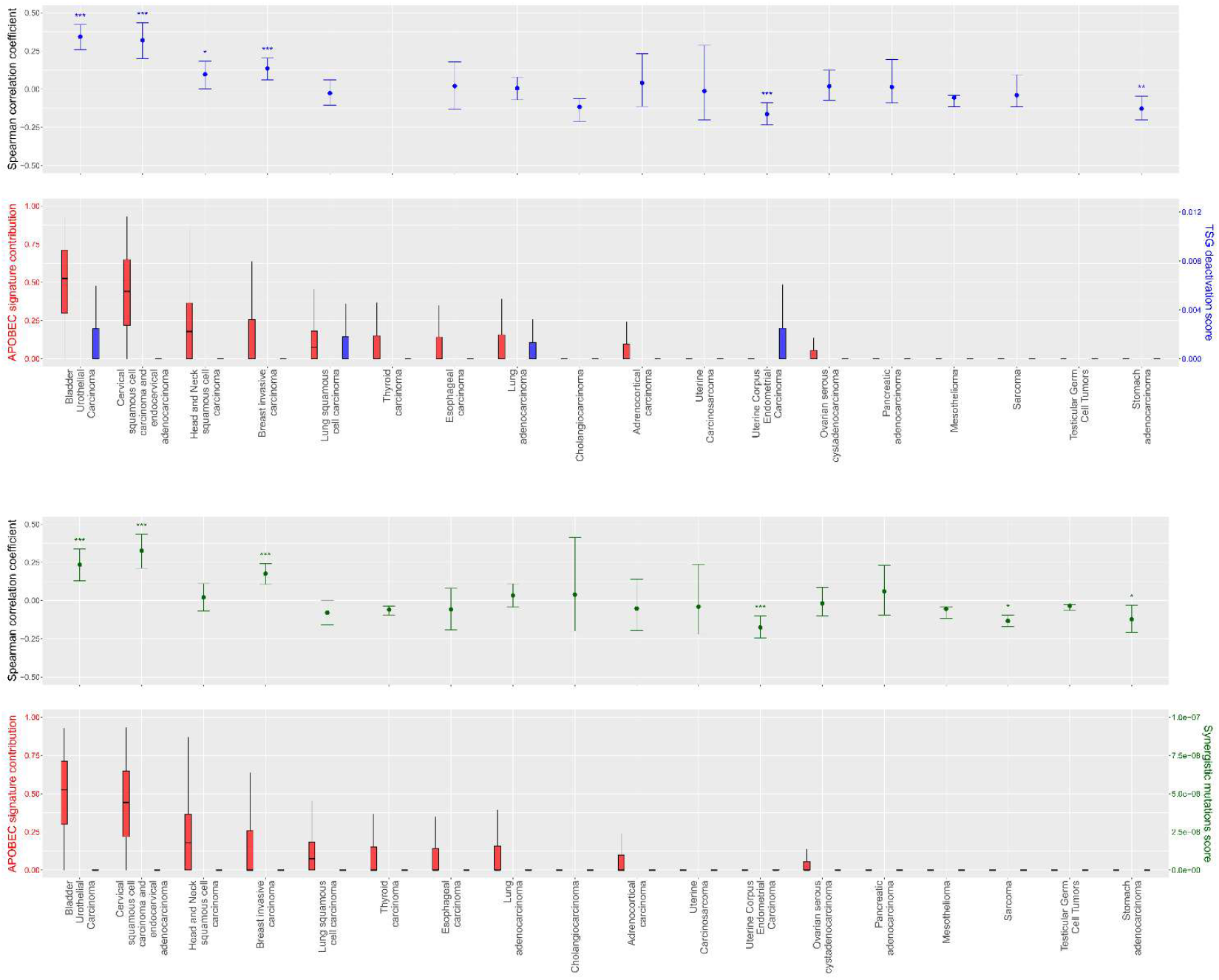
Robustness to choice of correlation coefficient. Distribution and Spearman correlation of APOBEC signature contribution and TSG deactivation score (top) or synergistic mutation scores (bottom), sorted by cancer type. Confidence intervals for Spearman correlation coefficients were computed using a percentile bootstrap with 1000 resamples at the 95% confidence level.

**Supplementary Figure 8:**
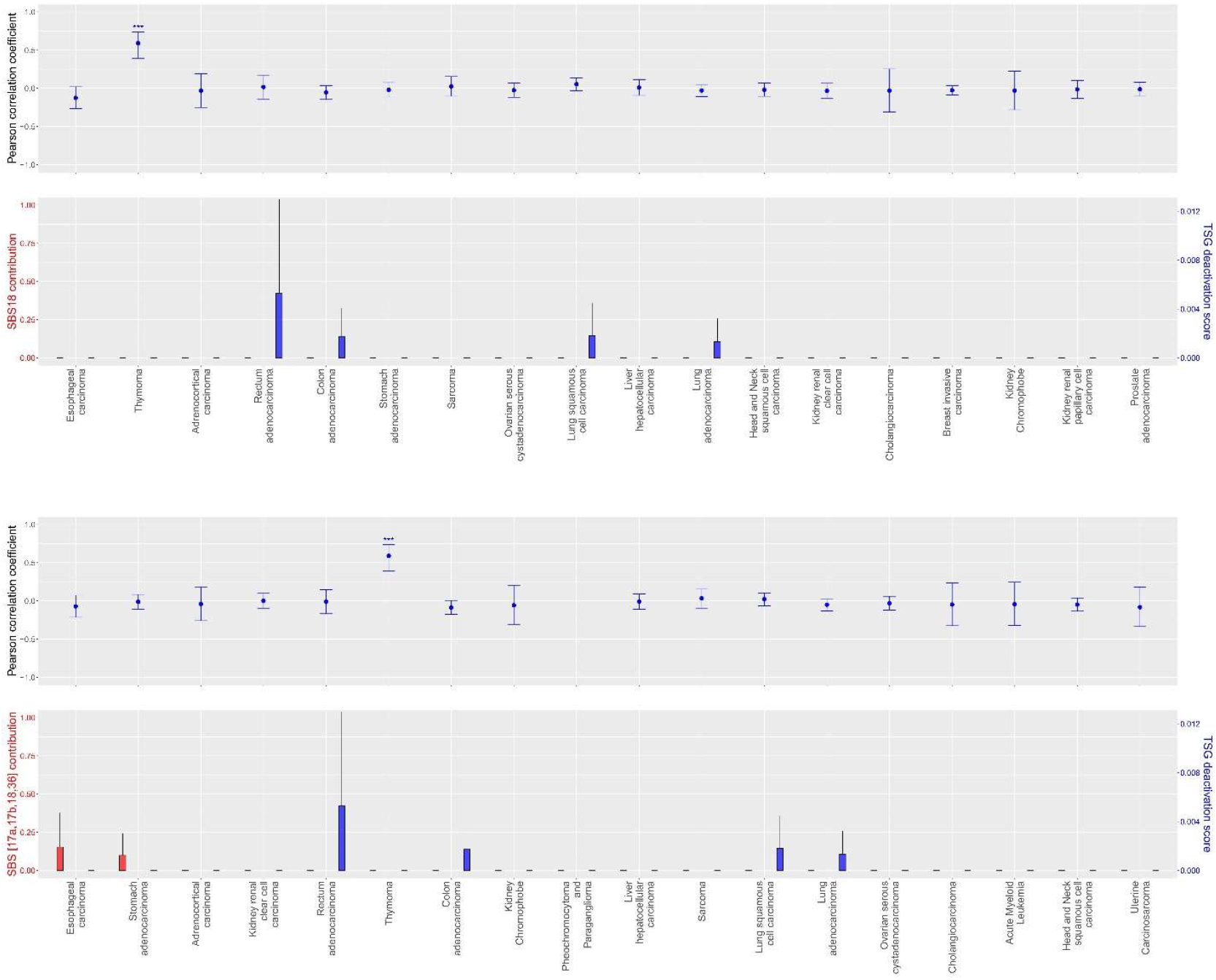
Robustness check: TSG deactivation and ROS in TCGA. Distribution and correlation of TSG deactivation score and ROS signature contribution measured as <u>SBS18</u> (top) or as the joint contribution of <u>SBS17a</u>, <u>SBS17b</u>, <u>SBS18</u> and <u>SBS36</u>. SBS18 only accounts for a small proportion of signature contributions and most correlations are close to zero. Including SBS17a, SBS17b and SBS36 which are less strongly associated with ROC does not change this correlation pattern relevantly.

**Supplementary Figure 9:**
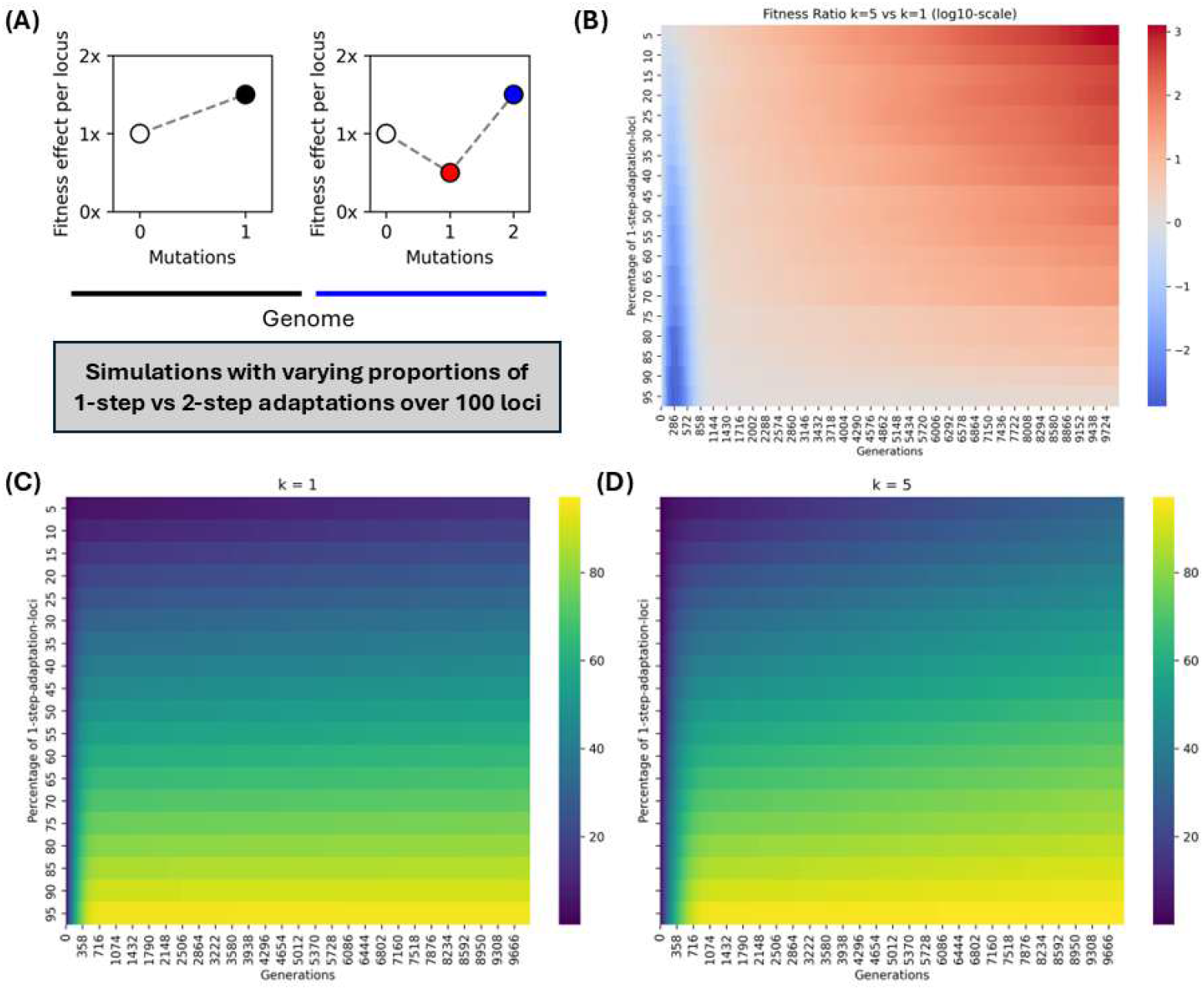
Temporal clustering and adaptation in fitness landscapes with varying ruggedness. **(A)** Sketch of the setup. Like in Fig 5, we assume that cells have two types of loci on which one-step or two-step adaptations are possible respectively. Fitness effects per locus are as in Fig 5. Here, we run simulations in which we vary the relative shares of the two types of loci. **(B)** Simulation results over time, averaged for 100 simulation replicates per row and clustering parameter k. Colors indicate the fitness ratio between simulations with k=5 and k=1 on a log10 scale. **(C)-(D)** show the corresponding absolute numbers of loci with adaptative mutations (i.e. loci in the black and blue states in panel (A)).

